# Hydrology-informed metapopulation modeling of liver fluke transmission in the Lawa Lake complex of northeast Thailand

**DOI:** 10.1101/569913

**Authors:** Tomás M. León, Vichian Plermkamon, Kittiwet Kuntiyawichai, Banchob Sripa, Robert C. Spear

## Abstract

While hydrologic processes are intuitively understood to influence transmission dynamics of water-related diseases, limited research exists that explicitly links hydrologic and infectious disease data. In the case of the life cycle of liver flukes, hydrology influences several transmission processes that mediate infection risk for multiple hosts. Northeast Thailand is a hotspot for liver fluke transmission and has strong seasonal flooding patterns. A metapopulation model linking local hydrologic processes with transmission of the liver fluke *Opisthorchis viverrini* in a lake system in northeast Thailand was developed and parameterized using infection data from 2008-2016. A rainfall-runoff model and other hydrologic data were used to assess level of connectivity between villages and the influence of upstream communities on parasite distribution in the study area. Disease transmission was modeled with metapopulations representing six village clusters around the lake using known prevalence data from humans, cats and dogs, snails, and fish. The metapopulation model improved upon the single-village model in its match to historical data patterns for the six village clusters with the introduction of the new time-variable parameters. Results suggest there are three unique hydrologic-epidemiologic regimes within the Lawa Lake system in response to upstream watersheds and risk of overland flooding that contribute to risk for *O. viverrini* infection. While available data may be insufficient to specifically characterize exact transmission dynamics, the practical implications of such findings are the importance of addressing connectivity for any intermediate host-based intervention. Similar approaches using hydrologic data to assess the impacts of water on pathogen transmission dynamics and inform mechanistic disease transmission models could be applied across other water-related disease systems.

## Introduction

Opisthorchiasis, infection with liver flukes of the genus *Opisthorchis*, is a disease whose transmission and distribution are significantly influenced by hydrologic processes. The parasites’ egg and cercarial forms require sufficient water and hydraulic transport to the next intermediate host for the transmission cycle to be sustained. Eggs are excreted in the feces of infected final hosts (humans, and reservoir cats, dogs, and other mammals to a limited extent); if not safely treated or contained, these eggs reach the aqueous habitat of intermediate host snails, which consume the eggs and mediate maturation to the cercarial stage. The cercariae are then released back into water, where they swim and seek out the second intermediate host cyprinid fish. They encyst in the fish, which if then consumed raw or undercooked by humans or other host mammals can migrate to their bile ducts and mature into adult worms. Water is a sustaining force for this parasitic life cycle, and its movement permits viable infection at each successive host stage.

The major liver fluke of interest in Thailand is *Opisthorchis viverrin*i. Given its known disease burden in Thailand, opisthorchiasis (infection with *O. viverrini)* has been a public health priority there, where it is transmitted to humans via the consumption of popular local raw and fermented fish dishes such as *koi pla* and *pla som*. The highest prevalence of opisthorchiasis and cholangiocarcinoma (CCA), the fatal bile duct cancer associated with *O. viverrini* infection, are found in northeast Thailand in the region surrounding Khon Kaen (Sithithaworn et al. 1997). Historical hotspots of opisthorchiasis and CCA include the villages around Lawa Lake. While much research has been conducted into the pathology of opisthorchiasis and CCA, there is limited literature addressing the ecologic and hydrologic aspects of parasite transmission in the environment (Grundy-Warr et al. 2012, Wang et al. 2017).

Lawa Lake is an approximately 4000-acre body of water that is highly vegetated and subject to significant hydrologic changes caused by seasonal variation in northeast Thailand. A peak in liver fluke infections is seen with lag following the rainy season in Thailand, as flooding facilitates the spread of fecal contamination and coincides with the rapid increase in snail populations (Sithithaworn et al. 1997). Since several weeks are required for the parasite to mature through its life stages, high infection rates in fish are seen in the late rainy season and late summer (July-January), and there is lower infection risk in the dry season and early summer (March-June). A primary industry on the lake is fishing, which contributes to sustaining local liver fluke transmission (Aunpromma et al. 2012). The hydrology of the Lawa Lake region is exceedingly complex and disturbed, as significant development in recent years has led to construction of new irrigation canals and ditches, new culverts and spillways that are opened and closed in the flooding season, and fish ponds that have become increasingly popular as a source of food and revenue.

Metapopulation modeling is commonly used to better understand the connectivity and influence of discrete populations and environmental patches on each other. This type of modeling is especially powerful for understanding pathogen transmission in heterogeneous environments for complex life cycles. Connectedness between these environmental and host patches can occur in multiple ways, including movement of humans between villages and dispersion of a waterborne pathogen from a section of river or lake adjacent to one population to another section adjacent to a separate population. The second example demonstrates hydrologic influence on a disease transmission system, as waterborne diseases as diverse as cholera and schistosomiasis rely on advective transport to expose new susceptible individuals with pathogens excreted or shed by infected individuals. Hydrologic patterns are time-varying and markedly local in nature. Given this environmental complexity, hydrologic fate and transport of pathogens are difficult to study. In addition, motile waterborne parasitic forms, such as liver flukes and schistosome cercariae, have independent mobility behaviors, making hydrologic flows not entirely representative of how these parasites reach their next host, though the distances covered are much smaller compared with the potential impact of flooding or river flow (Haas et al. 1990, Krishnamurthy et al. 2017).

Research connecting hydrology with waterborne disease transmission is an emerging field with recent work on cholera and schistosomiasis (Remais et al. 2008, Rinaldo et al. 2012, Perez-Saez et al. 2016). Tracing the spread of pathogens in the environment is challenging, and countervailing forces and feedback loops make it difficult to ascribe an increase or decrease in human infection to trends in meteorology and climate or consequent hydrology. Long-term studies of climatic changes in rainfall patterns or the influence of dams allow more definitive statements about impacts on disease transmission, though these are also complicated by shifts in host and vector habitat and in seasonal patterns that may disrupt or exacerbate host and vector growth and reproduction (Tompkins et al. 2013, Ziegler et al. 2013).

In this work, a metapopulation disease transmission model is developed and parameterized to assess hydrologic connectivity and *O. viverrini* parasite movement between six village clusters around Lawa Lake in Khon Kaen Province, Thailand, and how that is reflected by opisthorchiasis prevalence in hosts. Understanding liver fluke transmission in this seasonal, hydrologically connected environment with modeling can help define the scale of transmission processes and thereby optimize environmental control and treatment to have maximum impact on reducing disease transmission and prevalence in this setting and others.

## Materials and Methods

The model structure is an extension of the modeling framework presented in León et al. (2018). The six village clusters studied (Figure 1) are now connected in a mechanistic metapopulation framework to account for exchange of parasites and hosts between village clusters and their associated environments. This enables the model to include the influence of population-level factors, spatial heterogeneity, and degrees of connectedness between patches. This metapopulation model leverages information about hydrologic connectivity between village and host clusters to understand the movement of the liver fluke parasite’s various forms in the environment as mediated by water. To consider local hydrologic impacts on the liver fluke transmission cycle, five main factors are included: 1) egg inputs into the system from upstream watersheds; 2) egg inputs into the system from overland flooding; 3) snail and fish mobility due to hydrologic connectivity; 4) snail and fish available habitat; and 5) hotspots where infectious snails come into contact with susceptible fish. These factors are modeled monthly with seasonality to account for changing patterns throughout the year, i.e., they are modeled in a piecewise fashion and updated monthly.

**Figure 1:**
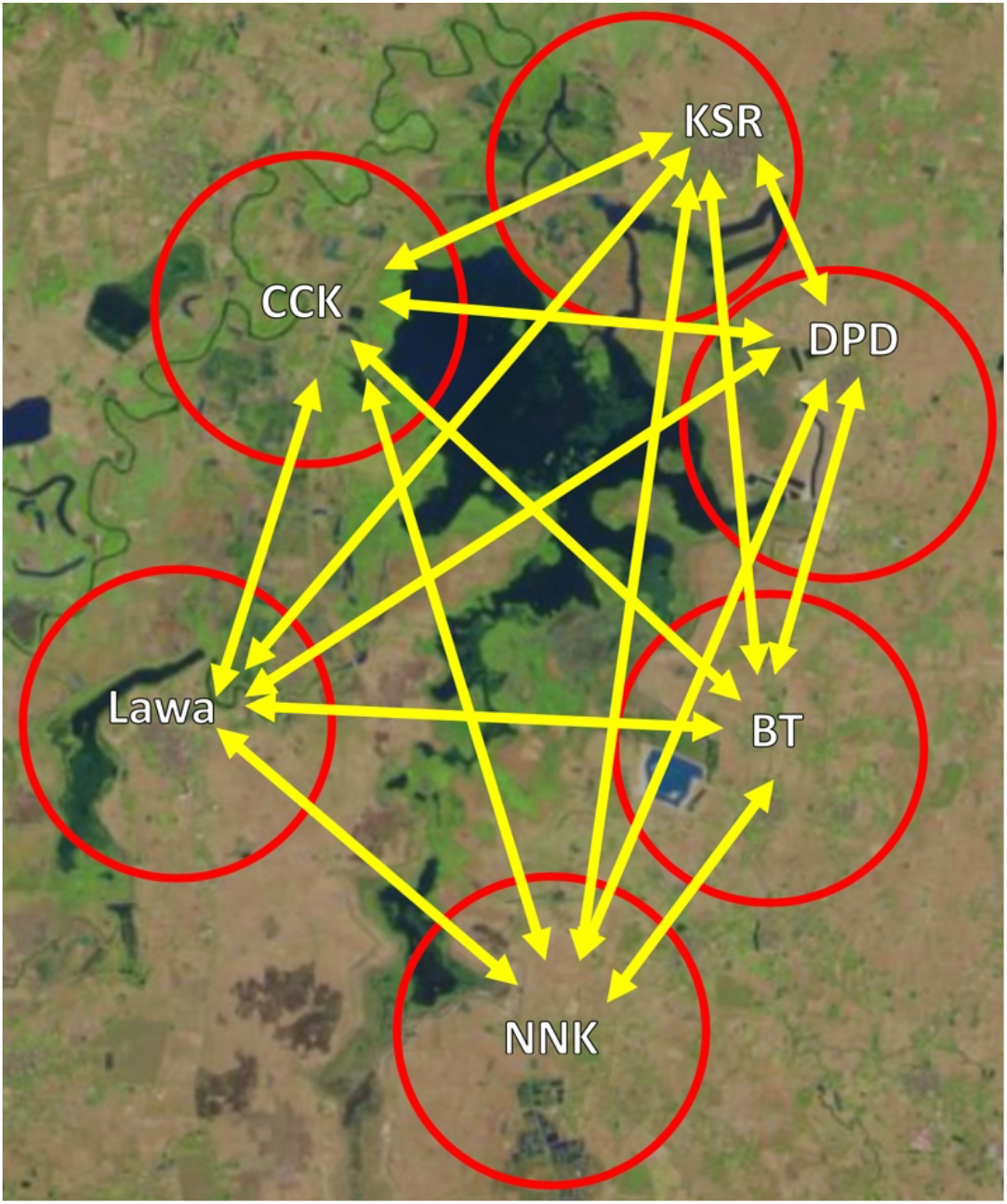
Connectivity between six village clusters around Lawa Lake (Map data: Landsat, USGS).

The metapopulation model connects the six villages or geographically proximate village clusters shown in Figure 1 around Lawa Lake in Khon Kaen Province of northeast Thailand. The villages or village clusters described here are CCK, Lawa, BT (cluster of 5 villages), NNK, KSR, and DPD (cluster of 2 villages), which were chosen and clustered based on geographical location and how historical human infection survey data was collected. The six clusters further sort into three groups based on impact or lack thereof of flooding and upstream watersheds. The two upstream watersheds (Figure 2) flow into Lawa Lake at locations adjacent to NNK and BT and contribute to egg input there. Villages CCK and Lawa are in proximity to the Chi River and are most susceptible to seasonal overland flooding. KSR and DPD are the villages most “downstream” and are relatively isolated from major flooding or upstream drainage impacts. Villages within the upstream watershed have not had as significant treatment and control programs as the villages around Lawa Lake, and infection surveys suggest that upstream villages still have high infection prevalence values over 50% (unpublished data).

**Figure 2:**
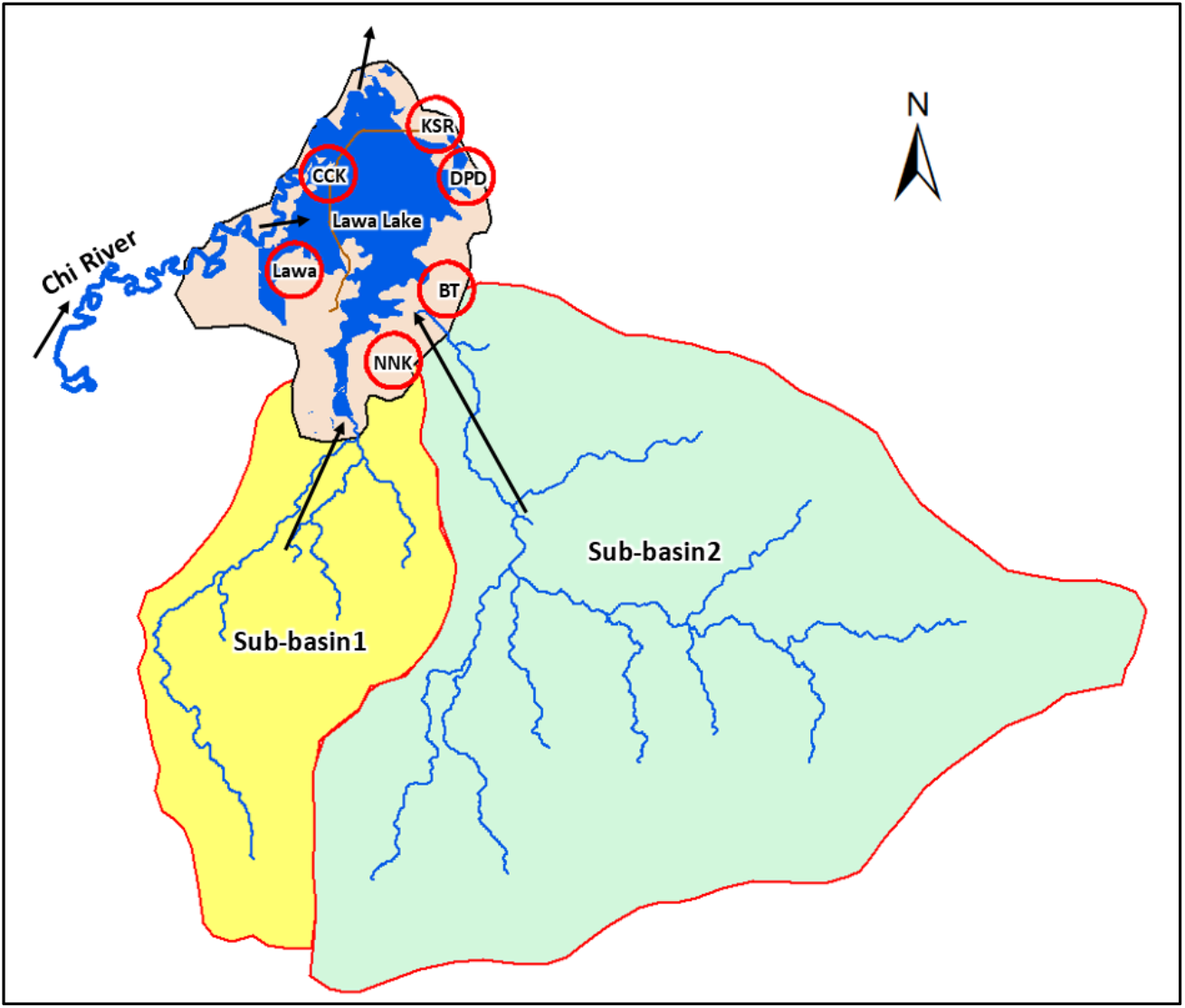
Two upstream sub-basins flow into Lawa Lake near NNK and BT, respectively (QGIS).

These hydrologic impacts seem to mirror trends in epidemiological patterns. Previous research highlighted the north-to-south gradient of nitrogen and salinity in the water that predicted higher snail abundance towards the south side of the lake (Kim et al. 2016). NNK, the southernmost village cluster, had the highest human infection prevalence at the time points when it was studied (Table 1). KSR and DPD, the northernmost villages and the farthest from the Chi River, had the lowest baseline prevalence of *O. viverrini* infection before the control program started. In the single-village model, varying transmission parameters by village did not fully capture the different patterns that occurred in the six village clusters when connectivity was not included. Therefore, the role of hydrologic processes and connectivity between villages needs to be considered in the disease transmission model to better account for the patterns observed.

**Table 1:** Infection prevalence (%) and mean intensity in positive individuals (EPG) for six village clusters around Lawa Lake. * indicates less sensitive diagnostic method (Kato-Katz or Kato thick smear).

| Village Cluster | 2008 | 2010 | 2011 | 2012 | 2014 | 2015 |
| --- | --- | --- | --- | --- | --- | --- |
| CCK | 54.9% (593) |  | 33.0%* | 44.3% (139) |  |  |
| Lawa | 67.1% (501) | 63.1% (108) | 19.0%* | (16) |  | 8.7%* |
| BT | 61.9% (346) | 37.2% (131) |  | 35.0% (136) | 9.0%* | 14.2%* |
| NNK | 74.1% (499) |  |  | 50.0% (61) |  |  |
| KSR | 16.4% (101) |  |  |  |  |  |
| DPD | 22.1% (112) | 36.5% (82) |  |  |  | 14.6%* |

To understand the effects of complex hydrologic factors on snail habitat, fish access to these habitats, and the pathways of parasite transmission, a hydrologic model of the Lawa Lake system was utilized to simulate flow patterns and changes in water levels over time. This model is a rainfall/runoff model common in hydrology that considers the transport of water through a system originating from upstream in the catchment basin or from precipitation. It uses a Soil & Water Assessment Tool (SWAT) model to generate runoff for the PCSWMM hydraulic model to determine hydraulic parameters of Lawa Lake such as flood depth and extent, flow velocity, and travel time (CHI 2018). The inputs for the SWAT model include meteorological data (rainfall, temperature, relative humidity, and windspeed) from the Thai Meteorological Department, soil type, land use, and a digital elevation model (DEM) generated from satellite imagery. For the hydraulic model, hydrologic structures and key parts of Lawa Lake were surveyed for elevation at 1m x 1m resolution using a drone, and a 2D model integrating runoff, the improved DEM, and meteorological data from 2008 to 2016 was developed using PCSWMM. Outputs include water level and flow vectors for the time points modeled between 2008 and 2016. Calibration was conducted with available precipitation and gauge data in the area from the Bureau of Water Management and Hydrology, Royal Irrigation Department, in Thailand. Figure 3 demonstrates an example of the variation in flows predicted by the model over the course of a calendar year encompassing the rainy and dry seasons; arrow direction and thickness represent the relative change in flow vectors. All of the villages are adjacent to Lawa Lake, and inflows and outflows as well as the impacted snail and fish populations are dynamic and heterogeneously distributed. The rainy season is characterized by high and active flows that generally peak in October with flooding from the Chi River varying from year to year. The dry season has relatively little hydrologic activity establishing connectivity between village clusters.

**Figure 3:**
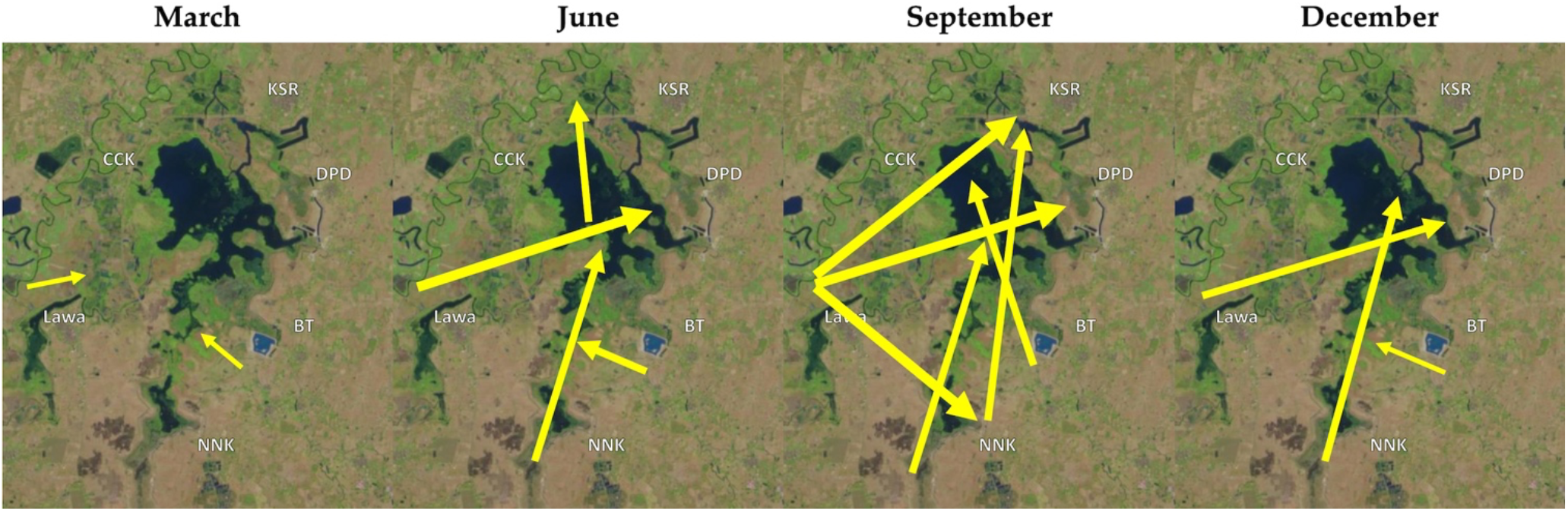
Hydrologic flows change dynamically throughout the year (semiquantitative interpretation of hydrology model results for an annual cycle) (Map data: Landsat, USGS).

Because gauge data was not available for the sub-basins upstream of Lawa Lake, a Soil and Water Assessment Tool – Calibration and Uncertainty Procedures (SWAT-CUP) model was used to calibrate and validate runoff into the Lawa Lake system from these sources generated by the SWAT model. Inputs for the upstream sub-basins included a 30m × 30m DEM, land use data, and soil type data from the Land Development Department of Thailand. To map the hydrologic features of Lawa Lake and finer scale structures, a drone was used to chart these areas in greater detail and determine elevations where water was flowing into or out of Lawa Lake. Sub-basin calibration and validation graphs are shown in Figure 4; 2006-2010 data was used for calibration, and 2011-2013 data was used for validation. The R^2^ values range from 0.61 to 0.81; both the calibration and validation models miss late peaks in their runs. For the calibration phase, fitting the less extreme peaks may have disadvantaged the model from predicting the major discharge in 2010. For the validation phase, the 2011 peak discharge was better modeled, but 2013 was missed by a large margin for reasons that are not entirely clear but may be related to the different timing of precipitation-driven flooding in 2013 compared with other years.

**Figure 4:**
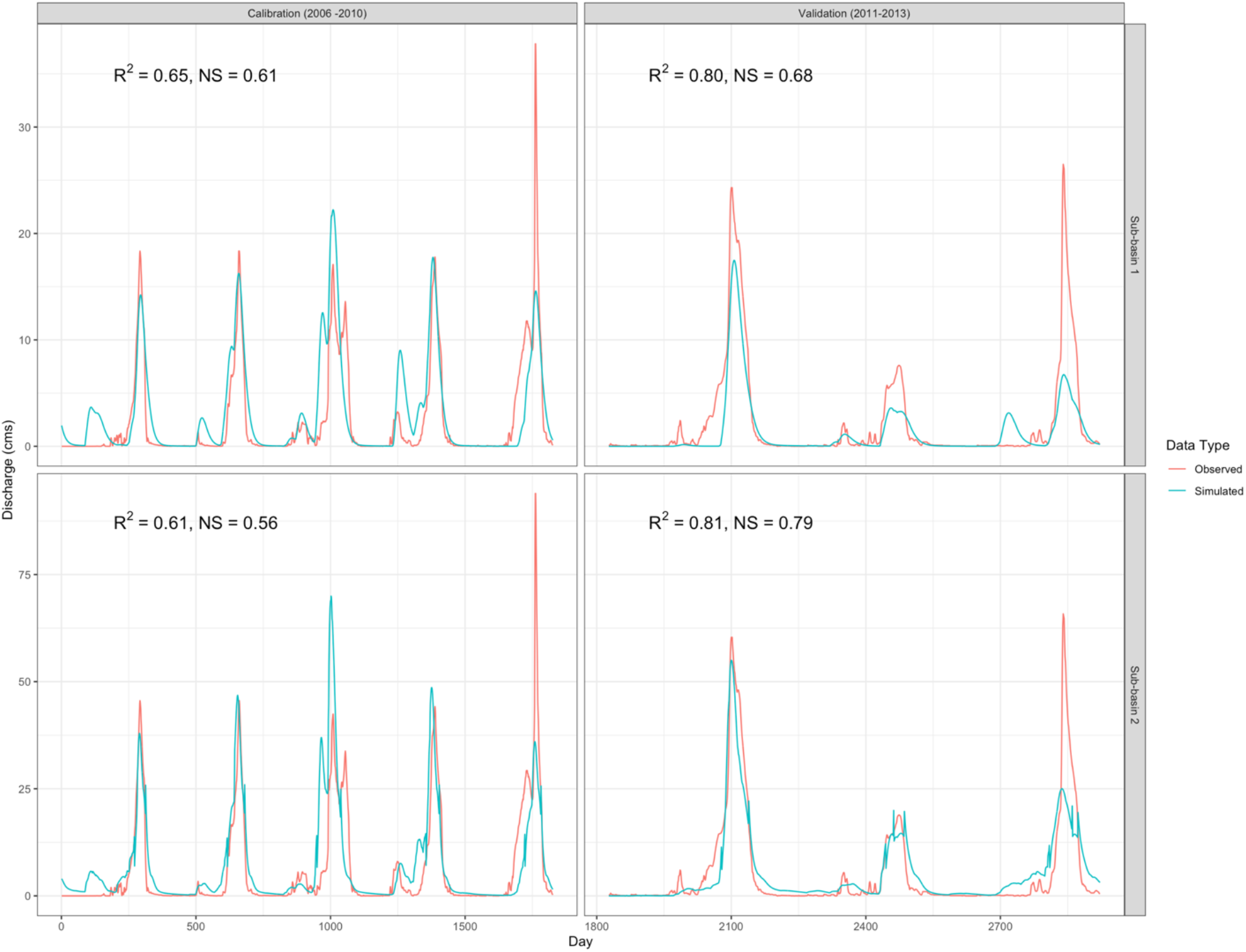
Calibration and validation curves for sub-basins 1 & 2. Calibration data encompasses 2006-2010 and validation 2011-2013.

Based on these data and tools, the presence/absence of connections between parcels of water associated with different village clusters were assessed. For example, KSR would not contribute to transmission in Lawa or NNK year-round but the reverse would be true. In March, when less rainfall and flooding occur, each village cluster is relatively isolated, with the exception of the relatively weak influence of near neighbors (Lawa to CCK or BT to DPD). The origin of flows is also subject to a differential dilution effect; contaminated waste with parasite eggs from the upstream watershed (Figure 2) would more strongly impact NNK than DPD or KSR, and the upstream villages (such as NNK) would experience the effects closer to discharge time.

The mathematical model of disease transmission (which incorporates data from the hydrologic model as variables and parameters) is an expansion of the single-village model described in León et al. (2018) to connect the six village clusters and uses as its state variables the infection prevalence in humans, reservoir hosts (cats and dogs), and snails, and infection counts in fish (equations in Appendix). Initial values were set from the baseline surveys in 2008, and base transmission parameters are carried over from the single-village model found using Markov chain Monte Carlo (MCMC) maximum likelihood methods to fit the model onto known infection prevalence data as described below. The infection prevalence data includes surveys using two different methods: formalin-ether concentration technique (FECT) and Kato-Katz (or Kato thick smear). In Thailand, FECT has been found to be the significantly more sensitive method to detect *O. viverrini* infection based on available data and because the protocol is intended to make microscopic examination easier (Laoprom et al. 2016, Saijuntha et al. 2018). Within the fish state variable is a fish demography model that captures the small window of time in the first few months of a fish’s life when it is susceptible to cercarial infection (before its scales harden and it becomes more resistant). This model assumes a maximum fish lifespan of 4 years before either being caught or natural death.

Figure 5 shows the general model flow diagram and how available data sources and village-specific factors are used in the model. The fecal waste-related egg inputs come from the Chi River and the two upstream watersheds and affect the snail state variable, contributing to the force of infection in that linkage; these time-varying parameters are derived on a monthly basis from the rainfall-runoff model. Other egg inputs from open defecation and disposal of septic tank sludge are not modeled due to lack of information about where and when they occur. The egg inputs from overland flooding of the Chi River were assumed to affect CCK and Lawa villages equally and were calculated by using flow measurements from the river and multiplying by a scalar to relate the impact of that water source with the upstream sub-basins. The first and second upstream sub-basins’ outflow were modeled to contribute eggs to the systems in BT and NNK exclusively and multiplied by their own scalars to translate those flows into contributions to human and reservoir host infection.

**Figure 5:**
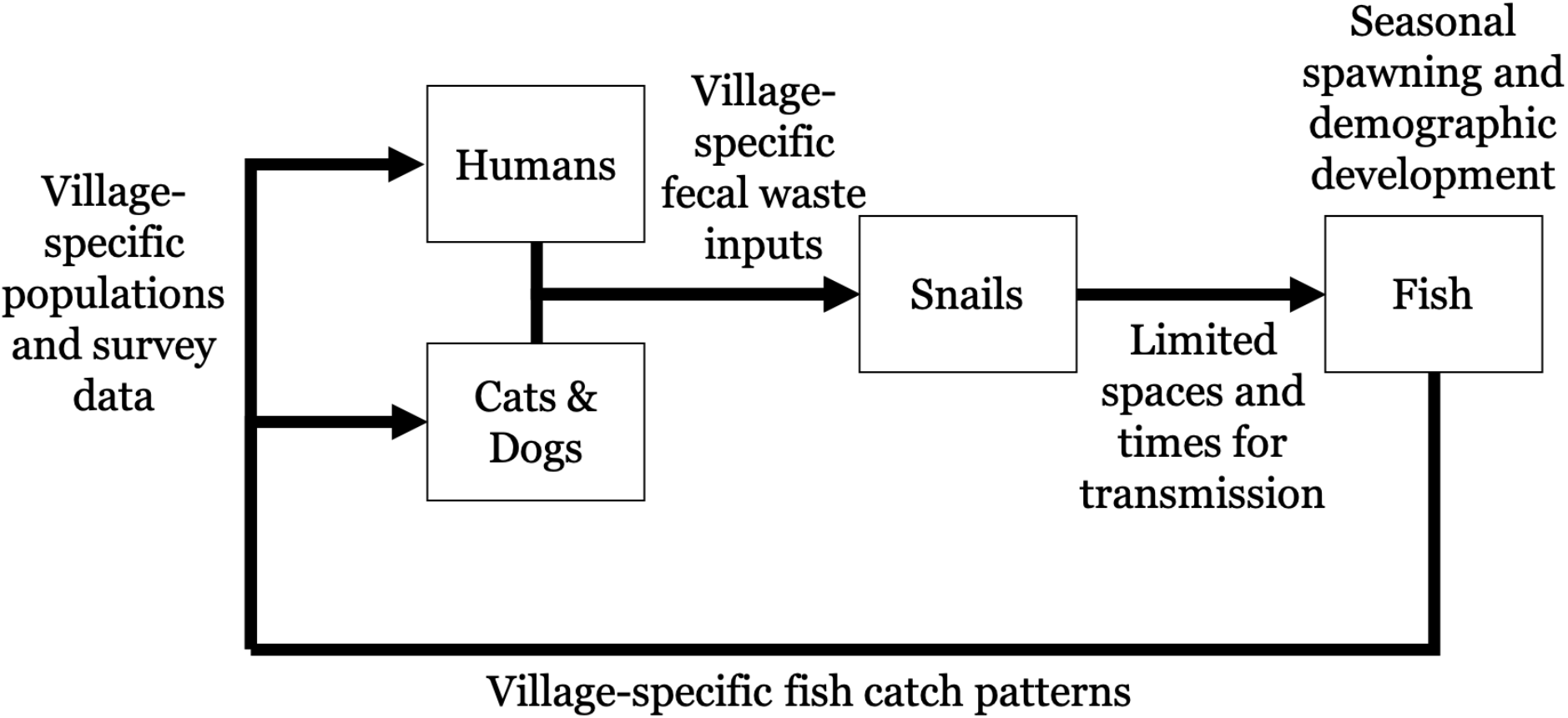
Model flow diagram showing submodels and input data that impact population numbers and transmission rates.

Connectivity rates between patches were varied on a monthly basis between 2008 and 2016 based on the hydrology model results to account for snail and fish mobility due to hydrologic connectivity. Figure 6 shows a model representation of how this works in a metapopulation model. These connectivity parameters were derived by assessing the fate and transport of parcels of water in a village cluster’s area and what proportions reached other village clusters in the Lawa Lake system. These ! parameters are unitless and vary from 0 to 1, describing the proportion of each village cluster’s force of infection for that host stage that affects each other village. Figure 7 shows examples of how these connectivity parameters varied by village pairing, month, and year (Figures 7–10 were produced in R using the ggplot2 package (Wickham 2016, R Core Team 2020)).

**Figure 6:**
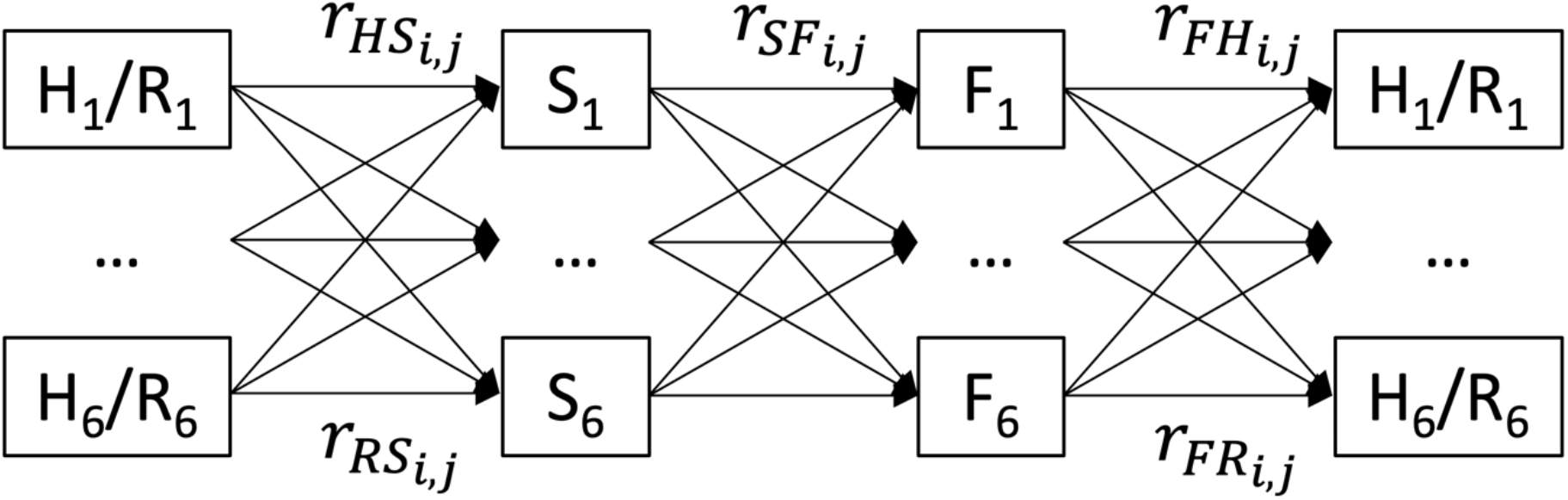
Model flow diagram showing how the metapopulation model works, with *r* connectivity parameters affecting transmission between each state variable. Subscript *i* denotes the source village and subscript *j* the receptor village for the parasite load in that transmission process. H = Human, R = Reservoir, S = Snail, F = Fish.

**Figure 7:**
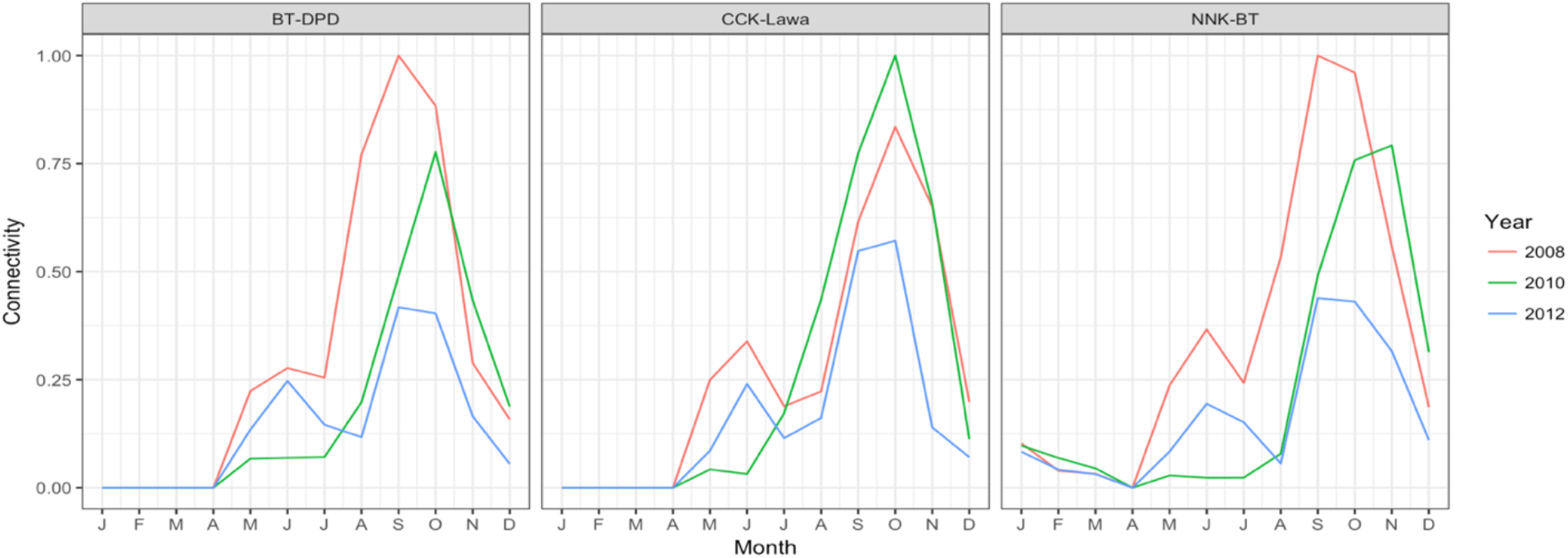
Village connectivity for BT-DPD, CCK-Lawa, and NNK-BT in 2008, 2012, and 2012

The parameters carried over from the single-village model are listed in Table S1. The β values are transmission parameters and are village cluster-specific (Table S2). The ! connectivity parameters are specific to each patch-to-patch relationship (Figure 1), γ is the fish catch rate describing the fraction of the total fish population caught at each time step, *λ*(*t*) is a gating function to control fish birth, death, and aging processes, *μ* are mortality rates, and α_*PZQ*_ are praziquantel (PZQ) treatment events.

While a daily time scale could be preferable for assessing hydrologic impact, historical data only captured month-to-month variability. Because human infection survey data only measures prevalence and not incidence, this time scale is reasonable for this study. From the hydrology model, the extent of water surface area at a suitable depth (under 0.3m) for the contact events between juvenile cyprinid fish and the aquatic snail intermediate hosts (“hotspots” for infection transfer) were used and to estimate snail populations N_S,i_ and fish populations N_F,i_. The transport time for a parcel of water between each village cluster was employed to estimate the time scale of movement between the locations, but these interactions happened on the order of days and not months and therefore the model’s time scale was not granular enough to introduce a time lag component. There was insufficient data to track fish mobility between patches, so fishermen’s movement data from Kim et al. (2017) was used to inform the exchange of fish in inter-village commerce as contributing to human infection from other village clusters. Table 2 summarizes these linkages and use of hydrology-related data in the disease transmission model. Figure 7 shows examples of the variability between village connectivity across months and years for the human/reservoir-to-snail and snail-to-fish transmission processes (the remaining connectivity parameter graphs are shown in the Appendix). The general trends persist from year to year, but the timing of peaks differ and affect village connections differently. The year 2008 produced stronger connectivity for BT-DPD and NNK-BT, while 2010 had stronger connectivity for CCK-Lawa.

**Table 2:** Description of linkages between hydrology model and disease transmission model

| **Hydrology-related transmission impact** | **Quantification method** | **Retained in model?** |
| --- | --- | --- |
| Egg inputs from overland flooding | Rainfall-runoff model output from Chi River summarized on monthly basis | Yes |
| Egg inputs from upstream water basins | Sub-basin model output summarized on monthly basis | Yes |
| Snail and fish mobility | Snail: Patch connectivity from rainfall-runoff model; Fish: fishermen catch data | Yes |
| Snail and fish available habitat | Snail: Rainfall-runoff model output and GIS analysis; Fish: N/A | Yes |
| Hotspots for snail-to-fish contact | Rainfall-runoff model output and GIS analysis | No |

## Results

Figure 8 shows the metapopulation model results for the six village clusters in the base scenario with the metapopulation model establishing relative connectivity between villages. Prevalence data points are included, distinguishing between the more sensitive FECT surveys and the less sensitive Kato method surveys. The heterogeneity of outcomes reflects the data: some villages saw reductions in infection prevalence to less than 10% (KSR, DPD), yet a few villages continued to have predicted prevalence values greater than 20% (BT and NNK). The steep drops in the graph were treatment events, when a subgroup of villagers was tested for infection and given praziquantel if they tested positive (the model assumes 100% drug efficacy). Model simulations were run for eight years between 2008 and 2016.

**Figure 8:**
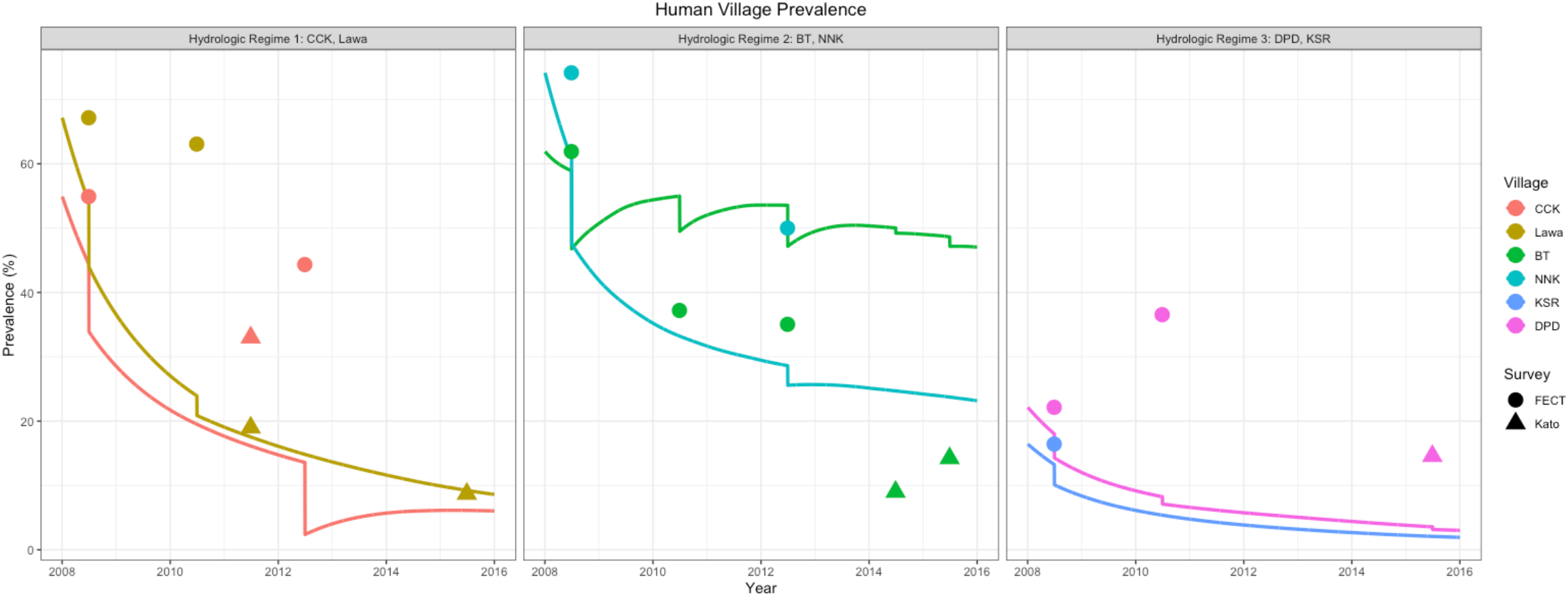
Metapopulation model run for human infection prevalence in villages around Lawa Lake

Figures 9 and 10 show infection prevalence values for intermediate snail and fish hosts. In snails, the prevalence cycles seasonally with most patch populations decreasing to 0.1% from initial values of 0.2% (with the exception of Lawa where prevalence approaches 0%). NNK has the highest final snail prevalence value at 0.18%, which is well within the range of what would be expected in this type of environment. For fish prevalence, because the initial conditions are disparate and based on baseline survey data, the model behavior is quite different. There is a seasonal aspect to their dynamics though this is dampened for most villages except NNK, where it is readily apparent. The end prevalence values range from 8-41%, with CCK, Lawa, BT, and DPD having the lowest values and NNK having the highest.

**Figure 9:**
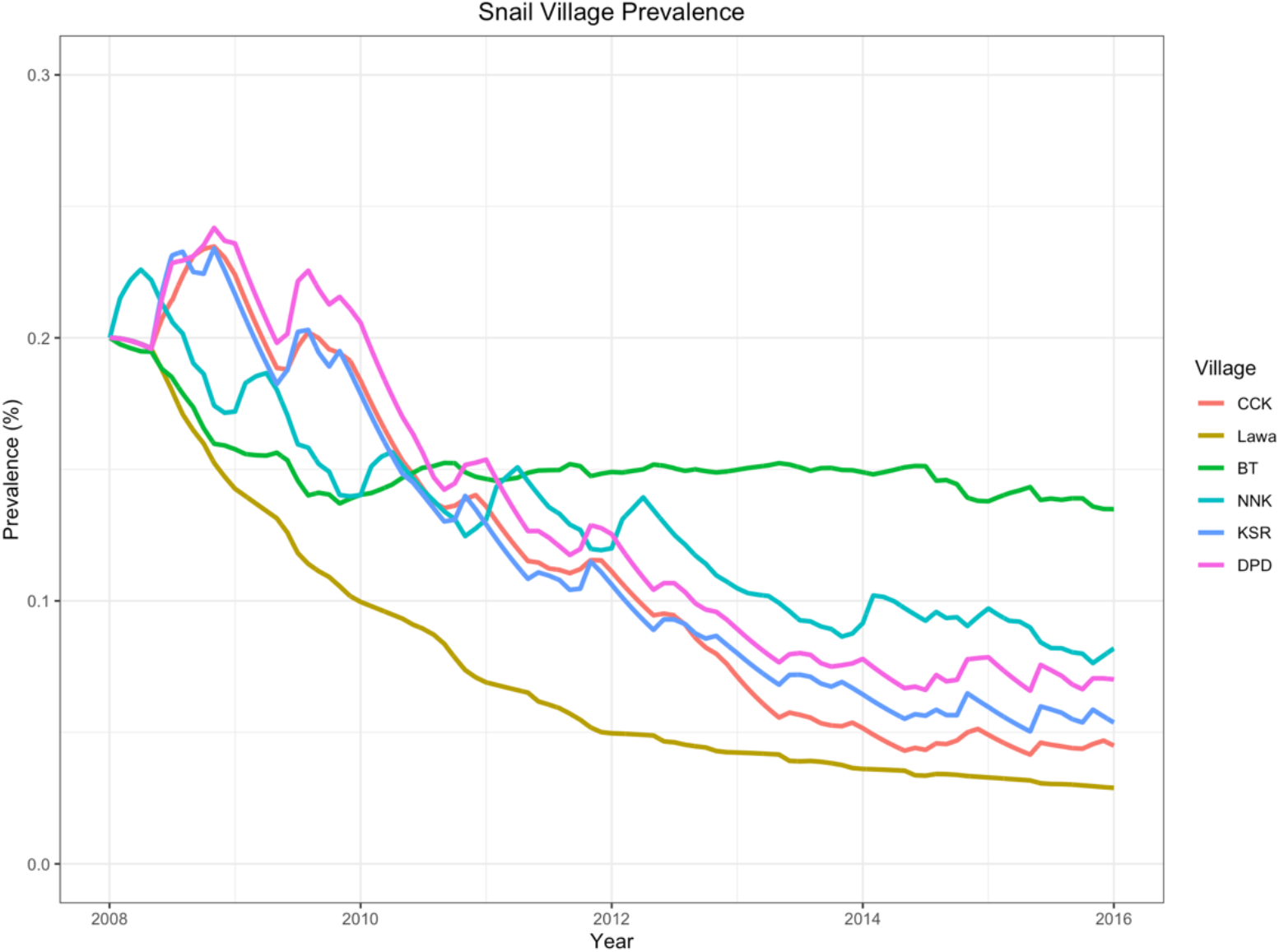
Snail prevalence values for metapopulation model

**Figure 10:**
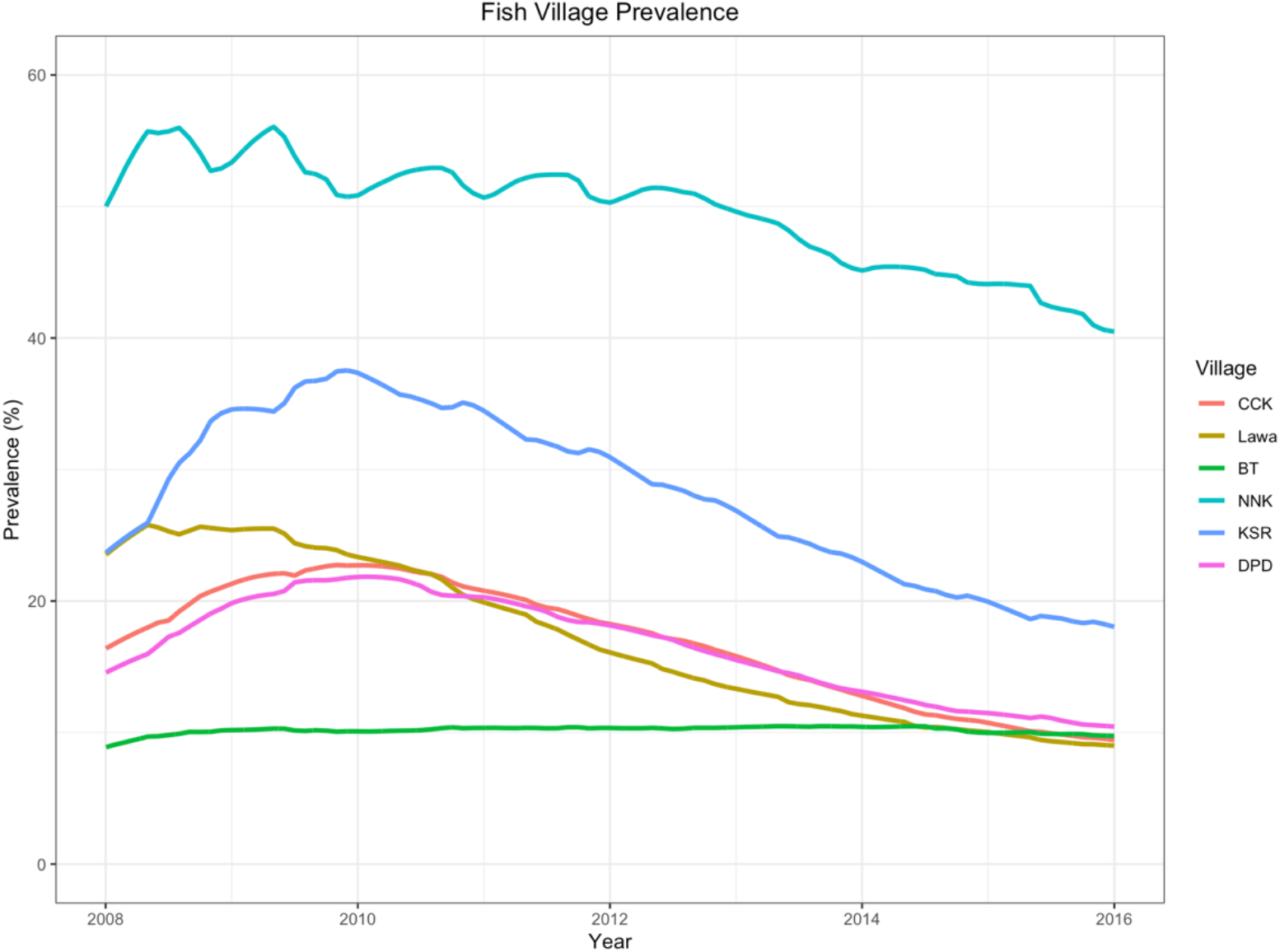
Fish prevalence values for metapopulation model

## Discussion

Compared with the single-village model, the metapopulation model no longer over-predicts final infection prevalence for the villages in 2016. Instead, the model now occasionally under-predicts prevalence for some data points, notably, CCK in 2012. This specific issue can be partially accounted for by the number of treatment-naïve individuals included in the 2012 infection surveys. Interpretation of the data leads to some specualtion about the meaning of the surveys and their different diagnostic methods. BT stands out as the modeled village with the least improvement (though it overpredicts the 2012 and 2014 data). According to control program managers, BT villagers were the least participatory in the Lawa Project and may therefore have reaped the least benefit from the control program. Given that this mathematical model is primarily concerned with infectious indviduals actively shedding parasites into the environment rather than asymptomatic cases, the Kato-based surveys from 2011, 2014, and 2015 may actually capture the most infectious and relevant individuals in the system and therefore be useful for thinking about infection prevalence patterns. However, fitting all of the data points accurately would be nearly impossible since most villages show non-monotonic patterns, infection burden builds up over time, and other unmeasured factors not included in the model may play a a major role in mediating these dynamics.

Two major questions are, given the discrepancy between model fits and the data, whether the data accurately reflects the reality of the disease transmission situation and whether the model should be tuned with yet more parameters to get a closer fit to the data. While this dataset is relatively complete and informative by the standards of NTD surveys, it still lacks enough time points, consistency in collection methods, and large enough sample proportions of the populations to give a detailed picture of the *O. viverrini* infection situation around Lawa Lake. The data (Table 1) show major swings across time points and discrepancies between the survey methods. Therefore, while the FECT data was used to fit via MCMC the transmission parameters in the single-village model, no parameters were fit for the metapopulation model because of the increase in model complexity and number of parameters (eighteen transmission parameters alone compared with three per single-village model, plus seventy-two time-varying connectivity parameters between the six village clusters). For this reason, the connectivity parameters were derived exclusively from hydrology submodel results. The metapopulation model is more believable than the single-village model in part because of its inclusion of external hydrologic influences and connecitivity and in part because the modeled behavior is more nuanced in the clustered patterns of village prevalence change it exhibits. The lack of parameters fitted to this model structure and the higher quality data that informed the hydrology model lend a realism to the underlying mechanics of the metapopulation model that improves upon the relatively straightforward transmission framework of the single-village model. The metapopulation model is meant to capture patterns of transmission rather than exact fits to the data. Nonetheless, the lack of information about differences in snail infection and raw fish eating patterns between villages remains a weakness, as they could not be incorporated into the model. Better data on these aspects of the transmission cycle would further strengthen the modeling framework and bring it into greater alignment with reality.

The patterns observed in these results support the sorting of the six village clusters into three geographical clusters that exhibit different trends based on human prevalence values. The first cluster, consisting of CCK and Lawa, is in close proximity to the Chi River and is most susceptible to overland flooding experienced during the rainy season. Its villages had high human prevalence values at the start of the control program, which decreased sharply during the period of treatment and control activity. These villages were the headquarters and major focal area of Lawa Project activities, suggesting that they benefited the most from health education and health volunteer engagement. The model is able to accurately account for the decrease in prevalence without making any assumptions about reinfection. Additionally, these villages are in close proximity to heavily fished waters in the lake, as supported by GPS evidence and interviews described in Kim et al. (2017).

The second cluster consists of BT and NNK, the villages to the south and southeast of the lake. These village also had high prevalence values at the start of the treatment and control program but experienced much more modest decreases when comparing data from later infection surveys. These villages are most impacted by upstream watersheds draining into Lawa Lake (Figure 2.2), where some villages still have over 50% *O. viverrini* infection prevalence (unpublished data). Consequently, if open defecation or unsafe disposal of human or reservoir waste is occurring in these watersheds, the runoff will disperse parasite eggs into the canals, ponds, and sections of the lake in close proximity to the second cluster’s villages. These villages are also adjacent to the highest concentration of fish ponds in the system and were not a focal area for Lawa Project activities.

The third cluster is KSR and DPD, which are located to the northeast of Lawa Lake. At the start of the control program, these villages had significantly lower prevalence values, which may be attributable to a lower degree of baseline environmental contamination. These villages were significantly affected by neither the Chi River nor the upstream watersheds, so they experienced fewer external inputs of infectious individuals or waterborne forms of the parasite into their local systems. These villages were not adjacent to high concentrations of fish ponds or fishing activity in their local waters and received less emphasis by the Lawa Project compared to the first cluster.

Because of the low prevalence of snail infection, the lack of field survey data, and the difficulty measuring snail prevalence precisely, strong claims cannot be made about the model results for the snail intermediate host. However, most field surveys indicate that snail prevalence in this region ranges between 0 and 0.2%, agreeing with the model results (Kim et al. 2016). With current diagnostic methods, differentiating between the clusters would require surveys of tens of thousands of snails at least. In recent years, the highest snail prevalence values found are still less than 10% (Kiatsopit et al. 2012). Much like other snail-borne diseases such as schistosomiasis though, only a few snails are required to maintain transmission in an area because of the high number of cercariae they shed into water bodies. Further understanding of where snails are most likely to be infected will help with environmental measurement and control. Bottlenecks of water flow, such as certain ditches and culverts, would concentrate fecal waste and parasite eggs and be zones of likely contact with susceptible snail hosts. Eliminating or protecting these areas could be an effective method of environmental control subject to proper coverage in the environment of interest and patch/cluster connectivity.

Fish infection prevalence is much higher than for snails and therefore it is easier to discern differences between clusters. Because transmission is foodborne, we are interested in the supply chain of food to consumers, which is not necessarily related to proximity between where fish lived and where they are eaten. Fishermen, middlemen, and merchants are all mobile and may choose to sell and distribute fish to other villages to expand their market. In the model simulation, the relative ordering of low to high fish prevalence values in fish hosts differ considerably from the results in humans, with NNK and KSR having the highest infection prevalence. NNK’s is driven largely by the initial value, but KSR’s is driven by dynamics, as its outcome is quite different from Lawa’s, which started with a similar prevalence level. Fish are infected by having infectious snails releasing cercariae into areas with juvenile fish, so KSR is the cluster with the greatest magnitude of this process taking place. Paying attention to fish prevalence results and how they interact with patterns of fish commerce can help identify where to target health education interventions related to cooking.

The model and the data that inform the model have limitations due to their fragmentary nature. Other model assumptions include ignoring the impact of different parasite burden levels in hosts and the age structure of human populations. The hydrology model was calibrated and validated against available data, leading to a plausible interpretation of the observed hydrologic behavior of the system. However, it could not account for very local effects that eluded its time and spatial scale and may have relevance for the points of contact between stages of the life cycle. The infection survey data may not be representative of the host populations because of sampling bias, but its overall spatial and temporal patterns align with local understanding and experience.

## Conclusions

This work highlights a major driver of persistent *O. viverrini* infection in northeast Thailand: a disturbed and dynamic hydrologic environment that mediates parasite transmission between connected village clusters and environments. This was accomplished by discussing and modeling five different means of hydrologic processes influencing parasite transmission and showing that its effects are significant and complex, acting heterogeneously across the Lawa Lake system. While local infection and contamination may be the main factor driving transmission at higher prevalence levels, as prevalence declines and villages move toward elimination connectivity will play a bigger role in maintaining the transmission cycle and preventing local elimination of the parasite.

The connectivity between water bodies and villages ensures that elimination of local infection is not possible without addressing upstream and adjacent environments. If infectious fecal waste from human and reservoir hosts is continually flushed downstream and the source is not treated, transmission will be restarted even if previously disrupted. This argument supports an approach that first targets villages and populations near headwaters and then proceeds further downstream while still accounting for human mobility and migration that could reintroduce infection into a previously cleared environment. Flood pulses and overland flooding also risk moving parasites into isolated and disconnected environmental patches on an annual basis, which requires constant treatment and attention to infection status of individuals in those patches. Snail and fish mobility remain little understood but have major relevance for *O. viverrini*’s life cycle, specifically how infection propagates in the environment. Targeting the locations where snails and juvenile fish come into close proximity with each other could be a promising environmental control technique but requires greater knowledge about the conditions that enable this transmission process.

A linked disease transmission-hydrologic modeling approach was employed here that uses hydrology model outputs as time-varying inputs in the disease transmission model to account for seasonal effects of flooding and water movement relevant to the intermediate hosts and waterborne forms of *O. viverrini*. Based on model results, village clusters were grouped into three disease prevalence curve patterns based on presence/absence of upstream and flooding impacts and history of control program intensiveness. Considering these findings, we argue for the use of this modeling approach and its results to inform environmental control of *O. viverrini* and for the need for environmental surveillance. While the specifics of the hydrology, population structure, and pathogen transmission cycle are local and specific in nature, this approach can be replicated across a variety of disease systems that are impacted by seasonality and a dynamic hydrologic regime.

## Acknowledgments

The authors would like to thank Barry Croke for helpful comments on the hydrology model and the Tropical Disease Research Laboratory staff at Khon Kaen University for their technical and logistical support for collecting the presented data.

The data used for this study are included in the main text and supplement. Analysis scripts are available from the corresponding author on reasonable request.

## Supporting information

**Table S1:** Parameter values for single-village model

| Parameter | Value | Units | Source | Symbol |
| --- | --- | --- | --- | --- |
| Natural mortality of snails | 1.37E-03 | per day | Kruatrachue et al. 1982 | $\mu_S$ |
| Parasite dependent mortality of snails | 1.37E-03 | per day | Chanawong & Waikagul 1991 | $\alpha_S$ |
| Mortality of fish | 6.85E-04 | per day | Suvarnaraksha et al. 2011 | $\mu_F$ |
| Parasite dependent mortality of fish | 0 | per day | Assumption (unstudied) | $\alpha_F$ |
| Natural mortality of humans | 3.69E-05 | per day | CIA 2015 (Factbook) | $\mu_H$ |
| Human infection clearance by praziquantel | Variable | Episodic | Treatment data from clinics | $\alpha_{PZQ}$ |
| Mortality of reservoir host | 2.74E-04 | per day | Local interview data | $\mu_R$ |
| Reservoir infection clearance by praziquantel | Variable | Episodic | Cat/dog treatment data from veterinarians | $\alpha_{PZQ,R}$ |
| Transmission parameters (fish-to-human, fish-to-reservoir, human-to-snail, snail-to-fish) | See Table 3.5.1 | per day per infectious host/worm | Equilibrium conditions and MCMC | $\beta_{FH}\beta_{FR}, \beta_{HS}, \beta_{SF}$ |
| Fish population | 3000 | fish | Estimate | $N_F$ |
| Snail population | 30000 | snails | Estimate | $N_S$ |
| Human population | Variable | humans | Village censuses | $N_H$ |
| Cat and dog population | 100 | reservoir hosts | Estimate from village censuses | $N_R$ |
| Mean human worm count | Variable | worms | Infection survey data | $W_H$ |
| Mean reservoir worm count | Variable | worms | Infection survey data | $W_R$ |

**Table S2:** Beta transmission parameters for single village model

| Equilibrium | CCK | Lawa | BT | NNK | KSR | DPD |
| --- | --- | --- | --- | --- | --- | --- |
| Fish to Human | 2.50E-10 | 7.24E-11 | 2.53E-10 | 1.81E-11 | 3.32E-07 | 2.68E-07 |
| Human to Snail | 8.90E-09 | 1.01E-08 | 4.92E-09 | 8.92E-09 | 6.56E-12 | 1.18E-11 |
| Snail to Fish | 7.34E-06 | 7.10E-06 | 3.40E-06 | 8.96E-06 | 1.67E-05 | 1.35E-05 |
| MCMC | CCK | Lawa | BT | NNK | KSR | DPD |
| Fish to Human | 2.95E-07 | 3.45E-07 | 7.26E-07 | 2.28E-07 | 3.30E-08 | 7.75E-08 |
| Human to Snail | 1.25E-08 | 2.53E-09 | 3.45E-09 | 2.26E-08 | 1.28E-08 | 1.48E-08 |
| Snail to Fish | 4.48E-06 | 7.03E-06 | 2.23E-06 | 2.28E-05 | 7.08E-06 | 3.89E-06 |

Model Equations

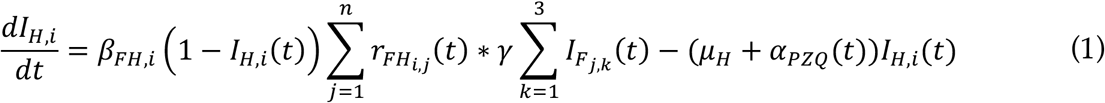

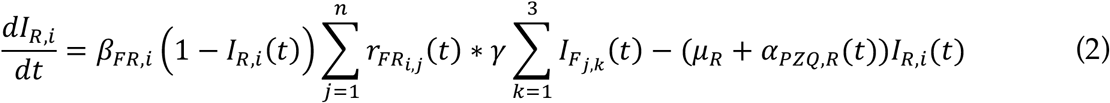

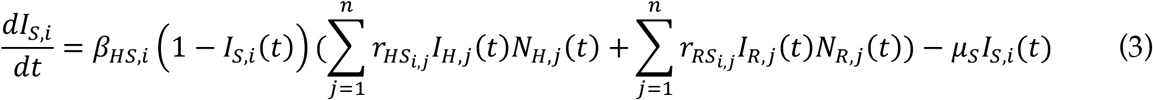

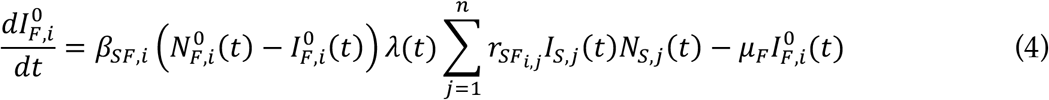

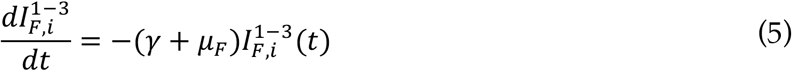

